# Long read and short read whole genome sequencing are equivalent for genomic characterisation of bacteriophage: considerations for high throughput analysis

**DOI:** 10.64898/2026.07.08.737226

**Authors:** Phoebe G Carr, Joshua J Iszatt, Mitchell G Hedges, Liza Mantjani, Andrew Vaitekenas, Stephen M Stick, Anthony Kicic, Samuel Montgomery, PhageWA

**Author notes:** Denotes corresponding author. Denotes co-first author (Phoebe G Carr, Joshua J Iszatt). Denotes co-senior author (Anthony Kicic, Samuel Montgomery).

## Abstract

**Background:** Antimicrobial resistance (AMR) is a global health crisis, necessitating alternative antibacterial strategies. Bacteriophages (phages) offer a promising solution, and their use as a therapeutic agent relies on stringent bioinformatic characterisation using whole genome sequencing (WGS) technologies. However, phages are highly diverse, with no clear consensus on best practices concerning phage DNA extraction or sequencing platform. Efficient and repeatable DNA extraction, sequencing, and bioinformatics processes are critical for safety assessments but remain poorly defined. Additionally, the impact of sequencing platform choice and DNA extraction methods on downstream genomic analyses is not well understood.

**Methods:** We evaluated multiple DNA extraction, library preparation, and sequencing approaches using a diverse collection of *Pseudomonas* phages from the PhageWA biobank. Column-based and precipitation-based DNA extraction methods were compared for DNA yield and recovery efficiency. Genome sequencing was performed using short-read (Illumina) and long-read (Oxford Nanopore Technologies) platforms, incorporating multiple library preparation kits and Nanopore basecalling models. Assemblies were assessed for completeness, quality, and sequence concordance using standardised bioinformatics pipelines, with hybrid Illumina-Nanopore assemblies used as references for comparison.

**Results:** DNA extraction efficiency varied substantially between protocols, with the Puregene precipitation-based method yielding significantly higher DNA recovery than column-based approaches when normalised to phage titre. Illumina sequencing consistently generated complete genome assemblies, although assembly fragmentation was observed for several jumbo phages when using the SeqWell ExpressPlex 2.0 library preparation method. For Nanopore sequencing, ligation-based native barcoding libraries produced longer reads than rapid barcoding libraries, while selection of the Dorado v5.0.0 basecalling model significantly improved read quality. Genome assembly success was dependent on phage genus; native Nanopore sequencing failed to assemble several Pbunavirus genomes, likely due to modified DNA bases, but an amplification-based library preparation successfully resolved these genomes. Across successfully assembled samples, Illumina and Nanopore platforms produced highly concordant genomes with comparable completeness scores, and hybrid polishing identified only minor sequence differences.

**Conclusions:** DNA extraction methodology, sequencing chemistry, and basecalling model selection significantly influence phage WGS outcomes. Precipitation-based DNA extraction improved DNA recovery, while both Illumina and Nanopore sequencing generated high-quality phage genomes suitable for therapeutic characterisation. Nanopore sequencing provided assemblies comparable to Illumina with minimal benefit from hybrid polishing, supporting its routine use for phage genomics. These findings provide practical guidance for phage genome characterisation workflows and contribute to the development of standardised, regulatory-grade approaches for therapeutic phage assessment.

## Introduction

Antimicrobial resistance (AMR) is a pressing global health crisis associated with almost 5 million deaths in 2019 [1], driving the need for novel antibacterial strategies [2]. Bacteriophages (phages) present a promising alternative due to their specificity and ability to target antibiotic-resistant bacteria [3, 4]. However, their clinical application remains limited, partly due to the lack of specific guidelines governing the genomic characterisation and regulation of phages [5]. Unlike bacterial genomes, phage genomes within a single species exhibit high diversity, making standardisation difficult [6, 7].While there are guidelines for phage genome assembly and annotation pipelines, [8-11] there is no clear consensus on best practices for phage genome assembly, annotation, or safety assessments for therapeutic use. The ability to perform robust safety evaluations of phages for therapeutic use hinges on accurate genome assembly, yet the impact of sequencing platform choice and extraction methodology on downstream genomic data remains remain poorly defined. As a result, the impact of upstream processes on downstream bioinformatics analyses is loosely characterised, complicating efforts to develop regulatory standards for therapeutic phages.

While bioinformatics pipelines for phage genome analysis continue to evolve rapidly, empirical validation of these tools remains limited and unsuitable for large scale reuse and reproduction [12, 13]. The rate at which computational pipelines are being developed far exceeds the generation of experimental data necessary to verify their accuracy and reliability, a process that is mitigated by peer review and noted to be absent from genome publishing processes [14]. One fundamental issue is the variability in DNA extraction efficiency across different phages, a process that must be developed and optimised by phage teams for the purposes of reliable manufacture. This variation may significantly impact genome sequencing outcomes and requires further investigation.

In addition to extraction challenges, a lack of direct comparisons specific to phage have been performed between sequencing technologies. Most research groups favour a single sequencing platform or library preparation and are typically informed by bacterial sequencing comparisons or manuscripts assessing a single technology [15, 16]. This has led to gaps in our understanding of the relative phage specific advantages and limitations of each approach for phage genomics efforts. While long-read Oxford Nanopore sequencing has recently become increasingly popular for bacterial genome sequencing, its utility for phage genome assembly and analysis remains underexplored. Given the potential impact of sequencing technology on genome completeness, contamination detection, and structural variation analyses, a systematic evaluation of these methods is warranted for phage research.

To address these gaps, we leveraged a diverse collection of *Pseudomonas* phages from the PhageWA [17, 18] biobank to investigate DNA extraction efficiency and sequencing technology impact. We systematically compared different DNA extraction methods, including column-based and precipitation-based approaches [19, 20], to assess their influence on the quantity and quality of extracted DNA. We then examined how these extraction methods affected raw read generation across both Illumina and Nanopore sequencing platforms. For Illumina sequencing, we tested multiple library preparation protocols to evaluate their impact on final assembly outcomes. Using high-throughput bioinformatics pipelines, we analysed genome completeness, contamination levels and sequence variation to assess phage genome characteristics in a standardised manner. This study provides critical insights into the methodological factors influencing phage whole genome sequencing data, with the goal of informing best practices and relevant considerations for high throughput screening and manufacturing efforts.

## Methodology

### DNA extraction

DNA was extracted from bacteriophage lysate. Host bacterial DNA was removed by incubating with 1uL of DNAse and 1uL of RNAse, and three different methodologies were tested in this study. Two methods used the Qiagen DNeasy column-based kit: our groups’ “EPIC” method based on Jakociune’s method [19], and a modified Harvard method “H2A” [21]. The third method precipitated bacteriophage from solution using zinc chloride [22], and extracted DNA utilising the Puregene Tissue kit (Qiagen) [23]. DNA quantification was performed using either the NanoDrop 2000 or the Quant-iT 1x dsDNA HS assay on the QuBit 4 (ThermoFisher).

### Illumina library preparation & sequencing

All Illumina library preparation and sequencing were performed by the Australian Genome Research Facility. Library preparations of bacteriophage DNA were conducted using either the SeqWell ExpressPlex 2.0 library preparation kit or the Nextera XT and 150bp paired-end reads were sequenced using the Illumina NextSeq2000, NovaSeq6000 or NovaSeqX sequencers.

### Nanopore library preparation & sequencing

DNA was neither sheared nor size selected before library preparation. Long-read whole genome sequencing of bacteriophage DNA was performed using the Oxford Nanopore MinION Mk1D platform. Libraries were prepared using either the Native Barcoding Kit 24 V14 (SQK-NBD114.24; Oxford Nanopore Technologies, UK), Rapid Barcoding Kit 24 (SQK-RBK114.24; Oxford Nanopore Technologies, UK), or Rapid PCR Barcoding Kit 24 V14 (SQK-RPB114.24; Oxford Nanopore Technologies, UK) and sequenced using MinION 10.4.1 (FLO-MIN114) flow cells. Raw reads from bacteriophages were basecalled and demultiplexed using Dorado [24] v1.3.0 with the dna_r10.4.1_e8.2_400bps_sup@v4.3.0, dna_r10.4.1_e8.2_400bps_sup@v5.0.0, or dna_r10.4.1_e8.2_400bps_sup@v5.2.0 basecalling models.

### Illumina assembly

Raw Illumina reads were taxonomically classified with kraken2 [25] v2.1.3 using the standard-8 kraken2 database (Accessed 04/09/2024), and bacterial reads were removed using extract_kraken_reads.py [26]. Genome assembly and putative phage contig identification were performed using the Phanta [27] v.0.4 pipeline with default configurations. Briefly, Phanta v.0.4 performed adaptor trimming, deduplication, and normalization with BBTools [28] v38.18, followed by read quality assessment with FastQC [29] v0.12.1. Genome assembly was then carried out using SPAdes [30] v3.15.4, with subsequent read mapping performed by BBTools [28] v38.18. Putative phage contigs were assessed for quality using CheckV [31] v1.0.3 with the CheckV database v1.5. Circular contigs containing terminal overlapping repeat sequences generated during assembly were identified using APC.pl [32], which detects identical sequence overlaps at contig ends indicative of circular assemblies. Identified overlapping repeat regions were subsequently removed to generate the final phage genome assemblies. Identified genomes were annotated and reordered with Pharokka [33] v1.8.2 using the Pharokka database v1.8.0.

### Nanopore assembly

Nanopore reads generated using the dna_r10.4.1_e8.2_400bps_sup@v5.0.0 basecalling model, which produced the highest read quality, were used for downstream assembly. Raw reads were taxonomically classified with kraken2 [25] v2.1.3 using the standard-8 kraken2 database (Accessed 04/09/2024), and bacterial reads were removed using extract_kraken_reads.py [26]. Reads were filtered for quality (<Q10) and length (<1000bp) using Chopper [34] v0.11.0. Read quality was assessed with NanoPlot [34] v1.46.2 before genome assembly with Flye [35] (--nano-hq, --meta) v2.9.6 with subsequent read mapping performed by minimap2 [36] v2.30 and qualimap [37] v2.3. Putative phage contigs were evaluated for quality using CheckV [31] v1.0.3 with the CheckV database v1.5. Identified genomes were annotated and reordered with Pharokka [33] v1.8.2 using the Pharokka database v1.8.0.

### Hybrid genome assembly and comparison

Hybrid assemblies were generated to obtain the highest-quality phage genome assemblies by polishing long-read Nanopore assemblies with paired short-read Illumina data. Illumina raw reads were first subsampled to approximately 1000x coverage using rasusa [38] v2.1.0, and the resulting reads were used to polish putative Nanopore phage genomes with pypolca [39] v0.4.0. Putative phage contigs were assessed for completeness and quality using CheckV [31] v1.0.3 with the CheckV database v1.5. The resulting polished assemblies were then used as reference genomes for assembly comparison. Illumina-only and Nanopore-only assemblies were independently compared against their corresponding hybrid assemblies to assess sequence variation and identify potential assembly errors associated with each sequencing approach. Whole-genome assembly alignments were performed using minimap2 [36] v2.30 with the asm5 preset, optimised for closely related genome assemblies, and sequence variants including SNPs and indels were identified from the resulting alignments using paftools.js within minimap2 [36] v2.30.

## Results

### Precipitation-based DNA extraction and a modified column-based extraction are best for DNA extraction from bacteriophage

To assess differences in DNA extraction efficiency from bacteriophages, we assessed extraction with a silica column-based extraction kit (Qiagen DNEasy) using two previously published protocols against a salting-out precipitation kit (Qiagen Puregene) methodology optimised in-house using a single bacteriophage isolate (E79). DNA extraction was significantly more efficient with the Harvard and Puregene protocols compared to the EPIC protocol (p<0.0001; Figure 2A). When the Harvard and Puregene protocols were assessed across 11 diverse bacteriophage genera, the Puregene protocol resulted in more DNA extracted per microlitre of lysate input (p<0.01; Figure 2B). Importantly, the total amount of DNA recovered correlated with the titre of the bacteriophage in the input lysate (rho = 0.747, p<0.0001; Figure 2C), so we assessed efficiency of DNA extraction normalised to the viral titre of the input and the Puregene method was significantly more efficient than the Harvard method (p=0.0003; Figure 2D).

**Figure 1:**
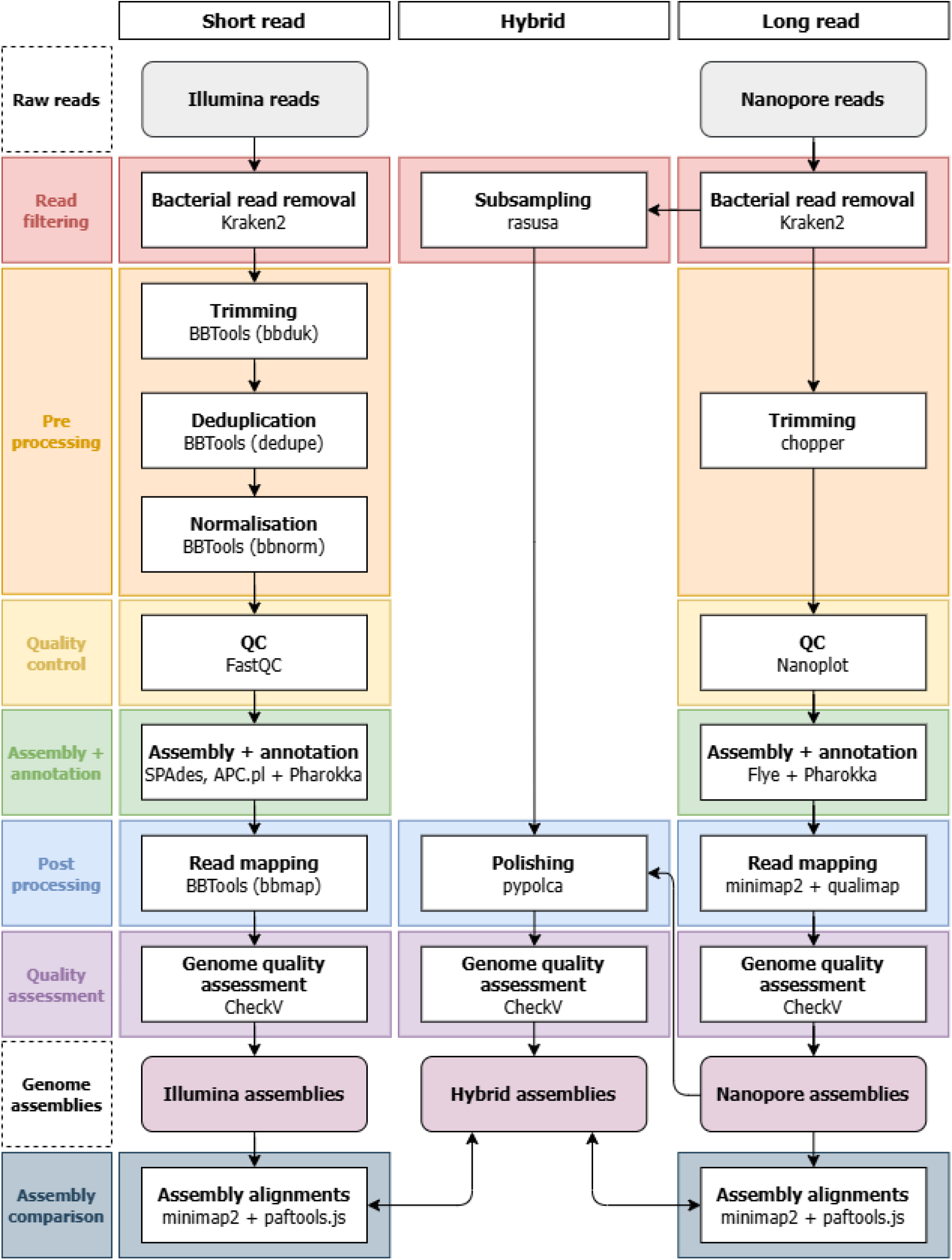
Genome assembly and analysis workflows for long and short read whole genome sequencing of bacteriophage.

**Figure 2:**
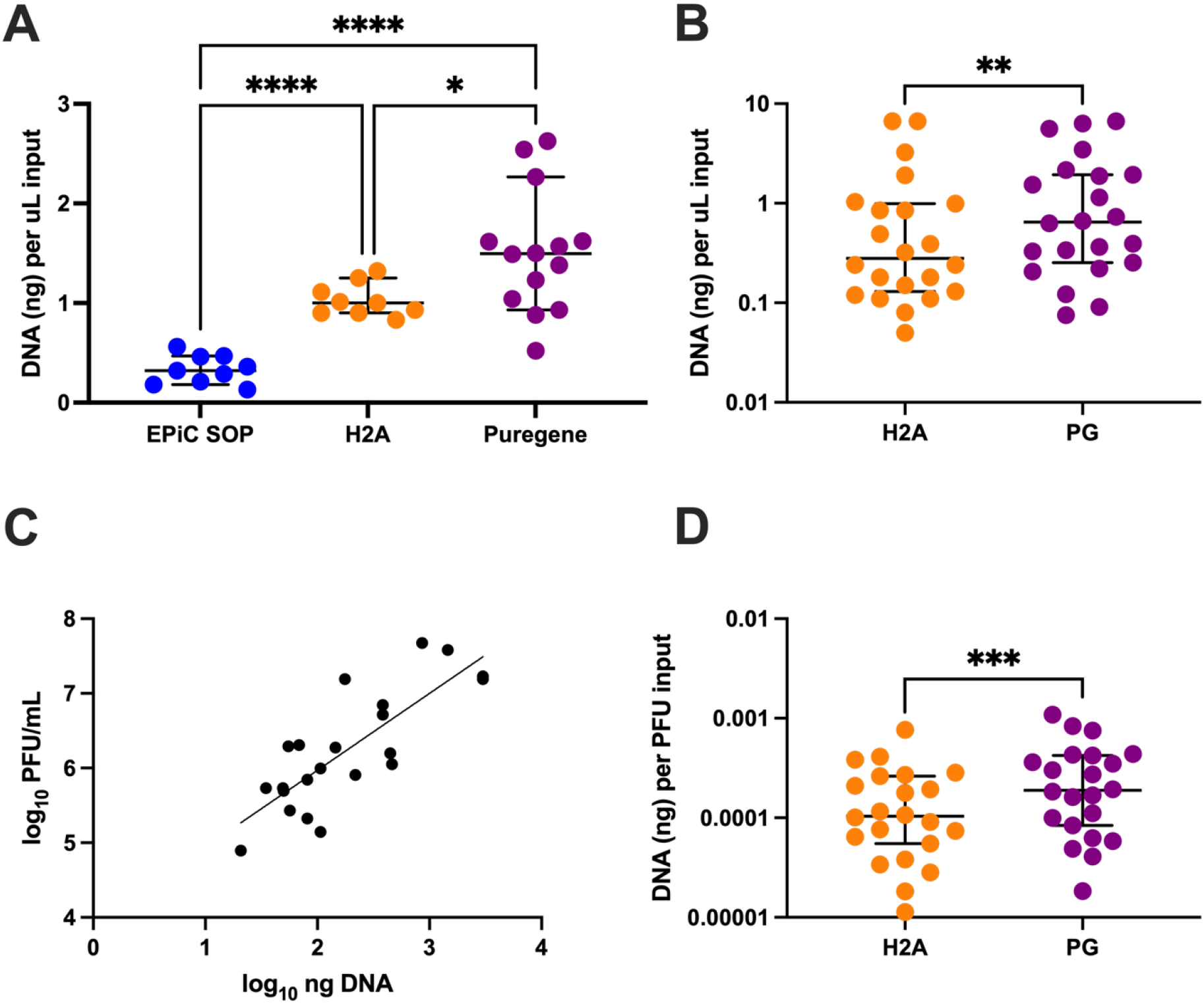
Optimisation of DNA extraction from bacteriophage lysate. DNA was extracted using a column-based kit using two previously published methods (EPIC, H2A) or a salting-out precipitation kit and method (PG). **(A)** Using a single bacteriophage lysate to compare extraction methods, the DNA recovered per uL of lysate using the H2A and Puregene methods was significantly higher than the EPIC method (n=9-14, p<0.0001; Dunnett’s T3 test). **(B)** When the H2A and Puregene were compared across diverse genera of bacteriophage, the DNA recovered from PG was significantly higher (n=22, p<0.01; Wilcoxon matched-pairs signed rank test). **(C)** The total amount of DNA recovered from both H2A and PG methods significantly correlated with the viral titre (PFU/mL) in the input lysate (rho=0.747, p<0.0001; Spearman correlation). **(D)** Normalised to the viral titre of the input lysate, DNA recovered from PG was significantly higher (n=22, p=0.0003; Wilcoxon matched-pairs signed rank test).

### Illumina short read sequencing enabled complete genome assemblies of bacteriophage

While short-read Illumina whole genome sequencing is considered the “gold standard” for viral sequencing, we assessed the impact of library preparation on bacteriophage genome assembly using two different library preparation methods. Phage samples (n=13) were prepared with both Nextera XT and SeqWell ExpressPlex 2.0 libraries. Reads were successfully generated from all samples and assemblies were produced using Phanta [27], a high-throughput assembly pipeline. Nextera XT libraries enabled single contig assembly of all phage genomes, whereas the SeqWell ExpressPlex 2.0 method yielded single contigs assemblies for only nine phages, and four fragmented assemblies. The four fragmented assemblies belonged to *Phikzvirus* (n=1) and *Pawinskivirus* (n=3), which were the only jumbophages within this sample set.

### Native Nanopore library preparation increases quality of data obtained through sequencing

To assess the effect of “tagmentaton” and ligation based Nanopore library preparation methods on bacteriophage sequencing, libraries were prepared from phages (n=10) using both Native Barcoding Kit 24 V14 (SQK-NBD114.24; Oxford Nanopore Technologies, UK) and Rapid Barcoding Kit 24 (SQK-RBK114.24; Oxford Nanopore Technologies, UK). Raw Nanopore signal data were basecalled using dna_r10.4.1_e8.2_400bps_sup@v5.0.0 dorado model and filtered for quality (>Q10) and length (>1000bp). There was a significant difference in both quality and read length between library preparation methods (p=0.0020, Figure 3A; p=0.0488, Figure 3B).

**Figure 3:**
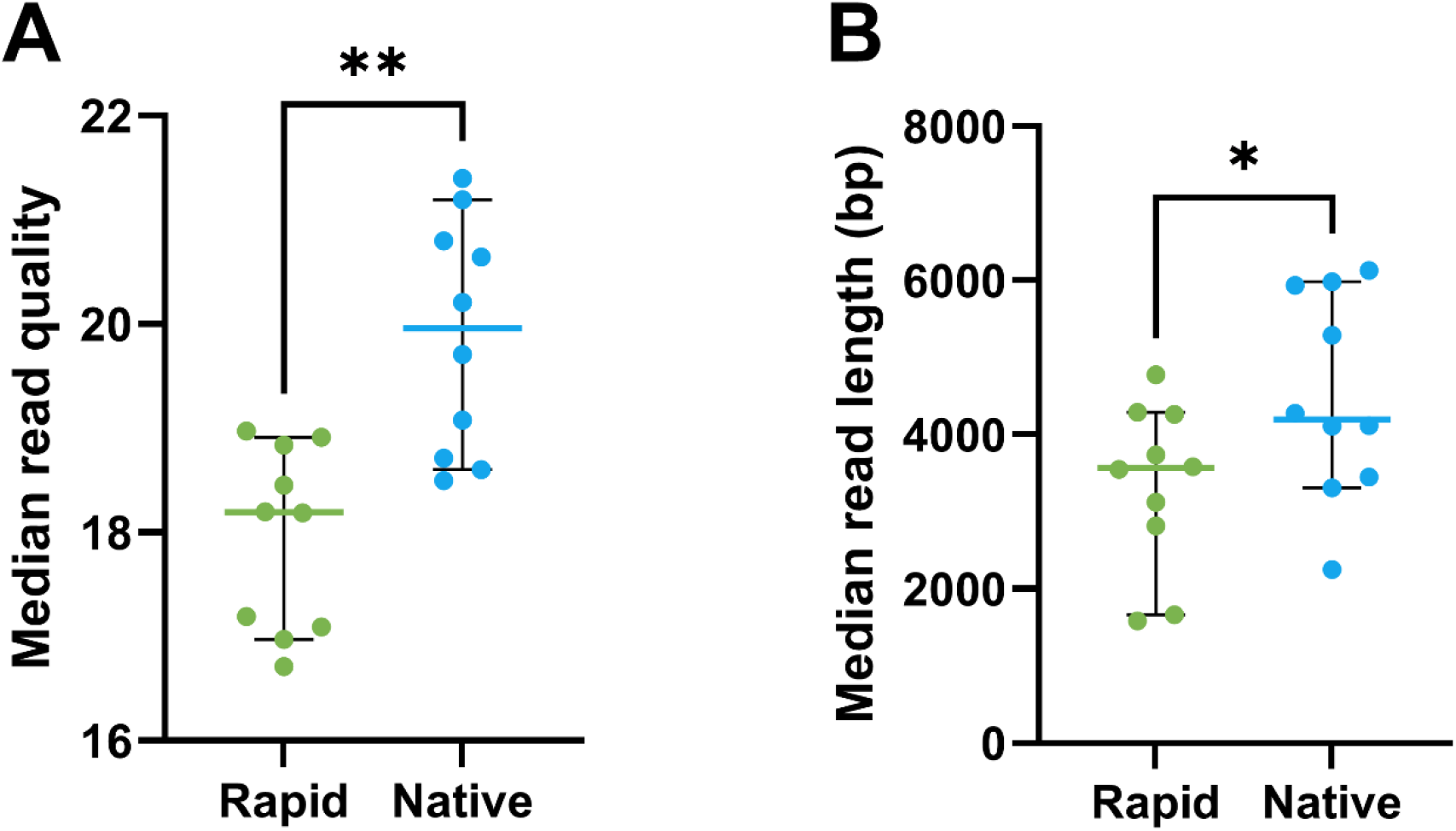
Ligation-based Nanopore library preparation increases read quality and length. When read metrics were compared across library preparation methods, ligation-based library preparation (Native Barcoding Kit) resulted in **(A)** a significantly higher median read quality (n=10 p=0.0020; Wilcoxon matched-pairs signed rank test) and **(B)** significantly higher median read length (n=10 p=0.0488, Wilcoxon matched pairs signed rank test).

### Nanopore basecalling model selection impacts read quality

As updated basecalling models were released by Oxford Nanopore during this analysis, we evaluated the impact of model choice on read quality. Raw Nanopore signal data from Native barcoded phages (n=47) were basecalled using the Dorado models dna_r10.4.1_e8.2_400bps_sup@v4.3.0, dna_r10.4.1_e8.2_400bps_sup@v5.0.0, and dna_r10.4.1_e8.2_400bps_sup@v5.2.0. After filtering reads (>Q10; length >1000 bp), choice of basecalling model had a significant effect on median read quality (Figure 4A). Notably, the v5.0.0 model produced higher-quality reads than both v4.3.0 and v5.2.0 and was therefore selected for downstream analysis. Following taxonomic classification, phage read sets were stratified by genus. *Pbunavirus* phages exhibited significantly lower read quality compared to all other present genera of phage present (Figure 4B), with all other genera of phage having similar median quality scores relative to other taxa (Figure 4C).

**Figure 4:**
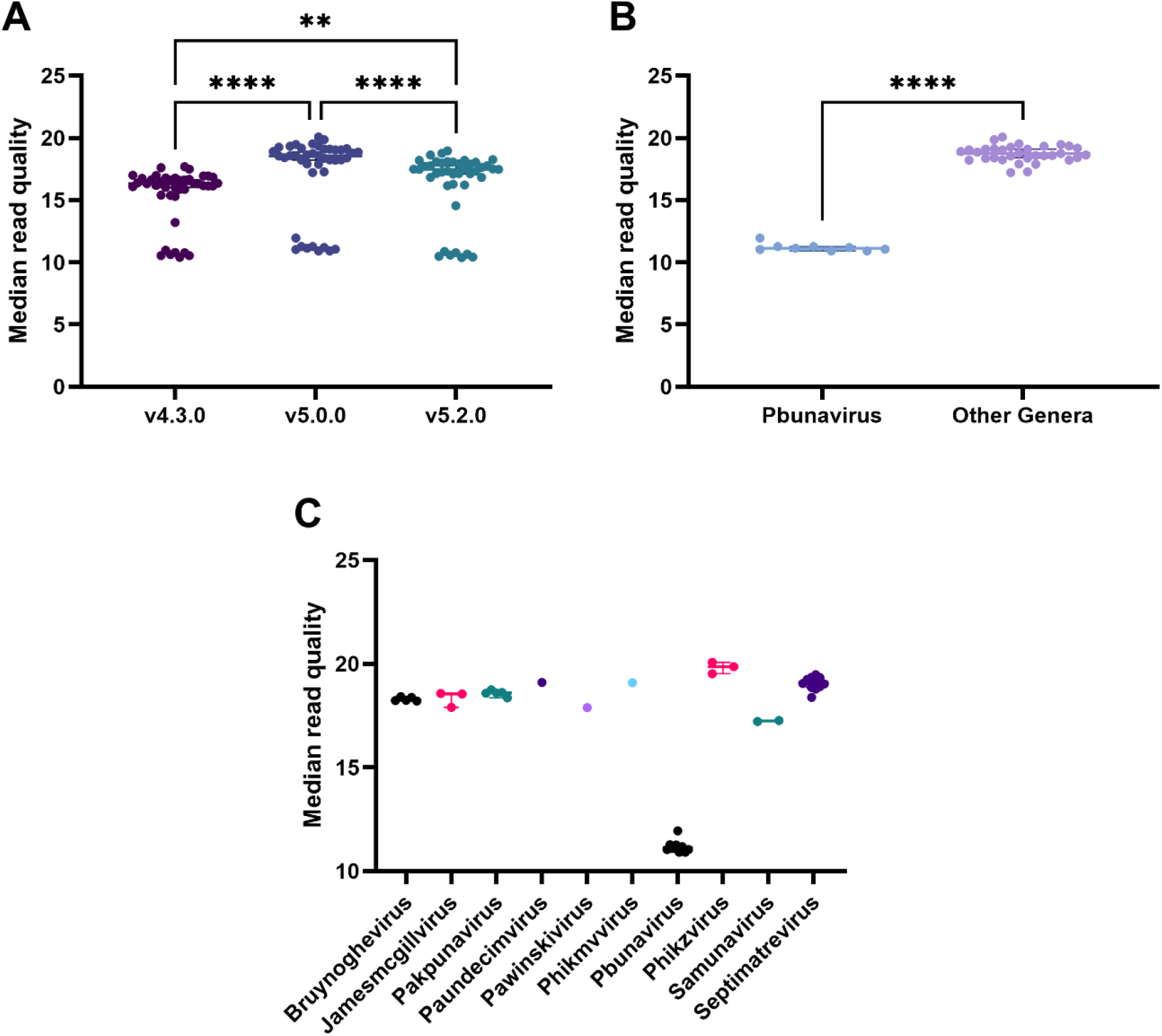
Basecalling model and bacteriophage genera have significant effects on read quality. **(A)** Quality- and length-filtered read metrics were compared across basecalling models. Reads basecalled with v5.0.0 exhibited a significantly higher median read quality relative to v4.3.0 and v5.2.0 (n=43, ****p=<0.0001 **p=0.0014; Friedman test). **(B)** Median read quality scores from the v5.0.0 basecalling model for *Pbunavirus* phages were compared to all other viral genera. *Pbunavirus* phages displayed significantly lower median read quality than reads from non-*Pbunavirus* phages (n=43, p=<0.0001; Mann–Whitney test). **(C)** Median read quality (v5.0.0 basecalling model) stratified by phage genus. Because most genera were represented by only one or two phages, formal statistical comparisons were not performed.

### Amplification-based Nanopore library preparation improves genome assembly for *Pbunavirus* phages

Of the 43 phages prepared using the Native Barcoding Kit 24 V14 (SQK-NBD114.24; Oxford Nanopore Technologies, UK), 33 were successfully assembled. The ten failed read sets were assessed for read quality and taxonomically classified, with comparison to corresponding complete Illumina assemblies confirming that all belonged to the *Pbunavirus* genus. Given the genus-specific reduction in read quality observed for *Pbunavirus* phages, an alternative library preparation approach was evaluated. Library preparation was performed for seven matching *Pbunavirus* phages using the Rapid PCR Barcoding Kit 24 V14 (SQK-RPB114.24; Oxford Nanopore Technologies, UK). Compared with the ligation-based native barcoding approach, rapid PCR barcoding produced higher-quality reads, with median read quality scores increasing from 11.26 (IQR 11.13–11.94) to 15.70 (IQR 15.60–15.80) across the seven phages (n=7, p=0.0006; paired t-test). Following downstream assembly, all seven phages were successfully assembled into complete genomes.

### Illumina and Nanopore sequencing generate similar bacteriophage genome assemblies

To directly compare Illumina and Nanopore sequencing for bacteriophage genome assembly, we analysed assemblies generated using Illumina SeqWell ExpressPlex 2.0 (n=36) and Nextera XT for jumbo phages (n=4), alongside Nanopore Native Barcoding (n=33) and Rapid PCR Barcoding for *Pbunavirus* phages (n=7). CheckV was used to assess the completeness of successful phage assemblies. Prior to removal of terminal repeat overlaps, several Illumina assemblies were identified as complete genomes with direct terminal repeats (DTRs), likely resulting from overlapping end sequences generated during short-read k-mer based assembly. Following removal of these assembly artefacts, all phage assemblies generated across both Illumina and Nanopore sequencing platforms were classified as “high-quality” by CheckV using AAI-based high-confidence completeness estimates. Completeness scores ranged from 94.55–100%, with no statistically significant difference observed between corresponding Illumina and Nanopore assemblies (p=0.59; Wilcoxon signed-rank test).

### Hybrid and single-platform assemblies show minimal genomic variation

To further assess assembly accuracy, Nanopore assemblies were polished using their corresponding Illumina reads to generate hybrid assemblies. These hybrid assemblies were then used as references for comparison against both Nanopore-only and Illumina-only assemblies. Comparison of Nanopore-only assemblies to the polished hybrid assemblies identified very few sequence differences. Detected variants were limited to isolated SNPs and single-base indels, with affected genomes differing by only one to two variants relative to their corresponding hybrid assemblies (Supplementary Table 1). All detected variants occurred within genes annotated as hypothetical proteins, except for phage BM83, where a variant was identified within a minor head protein gene. No substantial structural variation or large-scale assembly discrepancies were observed between Nanopore-only and polished hybrid assemblies. Comparison of Illumina-only assemblies to the polished hybrid assemblies identified sequence differences in only two phages.

One phage contained a single variant within a hypothetical protein that did not alter the resulting annotation. The second phage, BM83, contained a larger difference spanning 224 bp within a minor head protein locus. The Illumina-only assembly contained a single coding sequence in this region, whereas the polished hybrid assembly contained three coding sequences spanning the same genomic interval and collectively encoding an equivalent total length. Two of the three coding sequences were assigned to the same PHROG as the Illumina-derived annotation, while the third lacked a PHROG assignment. The corresponding minor head protein locus also contained a variant in the Nanopore-only assembly.

## Discussion

This study systematically evaluated how upstream factors, including DNA extraction method, library preparation, sequencing platform, and base-calling model influence the quality and completeness of phage whole genome sequencing outputs. By comparing column- and precipitation-based DNA extraction protocols across both Illumina and Nanopore platforms using a diverse panel of Pseudomonas phages, we provide insights that inform high-throughput phage genomic characterisation strategies and support the development of standardised practices for therapeutic phage validation.

By optimising input volumes and including chemical denaturants in our DNA extraction protocols, we significantly increased our yields of phage DNA whilst limiting the effects of bacterial host DNA interference. The significant improvement in phage DNA yield between our initial EPIC method and the optimised H2A and Puregene methods is attributed to the inclusion of the strong anionic detergent sodium dodecyl sulphate (SDS) which denatures proteins, such as those comprising the phage capsid [40]. SDS is contained within Buffer ATL in the column-based extraction, and the Cell Lysis Buffer in the Puregene kit. For our phages, SDS-proteinase K mediated denaturation of the phage capsids liberates greater amounts of phage DNA when compared to proteinase K alone, such as the EPIC method or Jakociune [19]. These results are consistent with other studies that employ SDS-proteinase K for phage capsid lysis [41, 42]. The significant increase in phage DNA yield resulted in high-quality sequencing and phage genome assembly.

Our findings demonstrate that the choice of basecalling model substantially affects both read quality and yield. Nanopore reads generated with the updated Dorado base-calling model (v5.0.0) consistently yielded higher quality data and significantly greater read numbers compared to the earlier v4.3.0 version (Figure 4A). This is consistent with other recent reports showing iterative improvements in Nanopore base-calling algorithms translating to tangible gains in output data quality. As basecalling software continues to evolve rapidly, keeping pace with these updates is essential for ensuring optimal performance in long-read phage sequencing. Despite this improvement, read quality varied across the phage genera tested (Figure 4B), with *Pbunaviruses* having significantly lower Q-scores across both base-calling versions (Figure 4C). This pattern suggests there is an underlying biological basis, likely due to modified nucleotides, that prevents specific nucleotides from being assigned a base label by the basecalling algorithm employed. Whilst the Nanopore basecalling software currently supports 5mC, 5hmC, 6mA, and 4mC detection, the modification type present in the *Pbunavirus* genomes investigated in this study may be causing the base caller to bin that base as “failed”, as it is unable to interpret the base signal.

This provides an explanation for the lower Q-scores for the *Pbunavirus* samples. An amplification-based library preparation method would overcome this limitation by producing non-modified DNA by whole genome amplification. Moreover, future updates to the basecalling software may include support for an expanded set of base modifications. Base modifications are increasingly recognised in phage biology as components that help phage escape the hosts’ CRISPR defence systems. It will become critical to understand and exploit such epigenetic modifications as bacteriophage become increasingly used as therapeutic agents in the fight against antimicrobial resistance [43]. This highlights a key consideration for the use of current long-read sequencing technologies, specifically ONT, in capturing the full diversity of phage genomes. Modified bases may currently present challenges for heavily modified phage genomes, however ONTs include a means to detect methylation patterns, and raw data can be re-basecalled to extract this information, providing advantages over short-read platforms. Furthermore, we demonstrate that the use of non-specific gDNA amplification via PCR may be used to remove base modifications and successfully sequence the phage genomes within this study.

Illumina sequencing, using SPAdes via the Phanta pipeline, reliably produced high-quality short-read assemblies, with 41 out of 45 samples yielding putative phage contigs. The remaining four initially fragmented assemblies were resolved through manual curation, highlighting both the robustness of short-read sequencing and the need for manual intervention for this data set. Nanopore sequencing produced 33 putative phage contigs from 45 samples, indicating a slightly lower success rate than Illumina. The failure rate was due to the sequencing of *Pbunavirus* phages, again suggesting lineage-specific challenges potentially linked to DNA modifications or repetitive elements that complicate long-read assembly.

Despite the slightly lower success rate, the Nanopore assemblies that were successful yielded contigs comparable in quality to Illumina, as assessed by CheckV completeness scores. This reinforces the use of long-read sequencing for generating full-length phage genomes. In our analyses, the use of either technology produced genomes of comparable quality. However, the inability of Nanopore native barcoding library preparations to resolve certain phages, particularly those belonging to the *Pbunavirus* genus, is a key consideration and underscores the continued relevance of short-read sequencing for comprehensive phage genomic analysis.

To evaluate whether combining technologies would produce more accurate assemblies, Nanopore-derived contigs were polished using Illumina reads using pypolca. The ‘hybrid’ approach led to minimal changes in the final assemblies, with a total of 13 errors (12 indels and one SNP) corrected across 29 genomes. This finding suggests that Nanopore sequencing alone, when sufficient in depth and read length, can produce phage assemblies of comparable quality to Illumina. This supports its use as a standalone sequencing platform for phage WGS. The limited benefits of polishing may be a result of the generally high quality and sequencing depth achieved for the raw Nanopore reads used in this study, particularly those generated with Dorado v5.0.0. However, the utility of hybrid assemblies may be context-dependent, and we caution against assuming universal benefit without prior evaluation.

Limitations within this study include that DNA extraction yield varied between phage types and it remains unknown whether all capsid types are susceptible to SDS-proteinase K lysis. Future work may include investigation of DNA extractions from phages belonging to different morphotypes to assess capsid susceptibility to lysis and suitability of extracted DNA for long read sequencing. Whilst our samples include a diverse array of *Pseudomonas* phages, broader generalisability to other genera (for example, *Staphylococcus, Acinetobacter, Burkholderia*) remains untested. Certain comparisons, such as the performance of different Illumina library preparation methods, were limited by batch, as library types were confounded between sequencing runs. While we attempted to control for this where possible, further head-to-head comparisons on matched samples would be valuable. Finally, while the CheckV tool remains a widely utilised standard for assessing viral genome completeness [31], its limitations include the influence of assembly artefacts [10] and its optimisation for metagenomic samples rather than isolate data.

Genome annotation across platforms was generally consistent with minor changes between Illumina, Nanopore, and Hybrid assemblies. Comparative analysis of coding sequences revealed no significant differences in key features related to phage activity, including genes associated with antimicrobial resistance, virulence, and lysogeny. This consistency supports the reliability of both sequencing platforms in identifying clinically relevant elements within phage genomes and reinforces their value for preclinical assessment pipelines. Future efforts should focus on functional validation of predicted genes, particularly integration-associated elements and tail fibre proteins that influence host specificity [44, 45].

This study assessed how DNA extraction methods, sequencing technologies, and basecalling models affect the quality and completeness of phage whole genome sequencing. Using a diverse panel of Pseudomonas phages from the PhageWA biobank, we show that Nanopore base-calling models significantly impact read quality and yield, with version 5.0.0 outperforming earlier versions. Nanopore assemblies were comparable to Illumina, with minimal benefit from hybrid polishing. However, certain phage genera, such as *Pbunavirus*, presented initial sequencing challenges linked to DNA modifications, but were successfully sequenced with an additional gDNA PCR amplification step prior to sequencing. The CheckV tool often misclassified DTR completeness in SPAdes assemblies, indicating a need for caution in automated assessments. Annotation results were consistent across platforms, reinforcing the suitability of either approach for therapeutic phage validation. These findings support the routine use of long-read sequencing in phage characterisation and highlight the need for continued tool development to support regulatory-grade genomic workflows.

## Supplementary Materials

**Supplementary Table 1.** Number of SNPs and indels identified in Nanopore-only and Illumina-only assemblies relative to their corresponding polished hybrid assemblies.

| Phage sample | Sequencing platform | Variants detected | Variant type(s) |
| --- | --- | --- | --- |
| 4VNF | Nanopore | 2 | 2 indels (1 bp each) |
| BM83 | Nanopore | 2 | 2 SNPs |
| J5TC | Nanopore | 1 | 1 indel (1 bp) |
| KMBS | Nanopore | 1 | 1 indel (1 bp) |
| U9X1 | Nanopore | 2 | 2 indels (1 bp each) |
| USCW | Nanopore | 1 | 1 SNP |
| UWKF | Nanopore | 1 | 1 indel (1 bp) |
| NWUG | Illumina | 2 | 1 SNP, 1 indel (13bp) |
| BM83 | Illumina | 1 | 1 indel (224bp) |

